# A Graph-Attention-Based Deep Learning Network for Predicting Biotech–Small-Molecule Drug Interactions

**DOI:** 10.1101/2025.05.13.653666

**Authors:** Fatemeh Nasiri, Mohsen Hooshmand

## Abstract

The increasing demand for effective drug combinations has made drug-drug interaction (DDI) prediction a critical task in modern pharmacology. While most existing research focuses on small-molecule drugs, the role of biotech drugs in complex disease treatments remains relatively unexplored. Biotech drugs, derived from biological sources, have unique molecular structures that differ significantly from those of small molecules, making their interactions more challenging to predict. This study introduces BSI-Net, a novel graph attention network-based deep learning framework that improves interaction prediction between biotech and small-molecule drugs. Experimental results demonstrate that BSI-Net outperforms existing methods in multi-class DDI prediction, achieving superior performance across various evaluation types, including micro, macro, and weighted assessments. These findings highlight the potential of deep learning and graph-based models in uncovering novel interactions between biotech and small-molecule drugs, paving the way for more effective combination therapies in drug discovery.

## Introduction

The increasing prevalence of complex diseases, particularly among the elderly, often necessitates the use of multiple medications. While polypharmacy aims to address different aspects of disease management and can lead to synergistic effects (e.g., aspirin for blood thinning with clopidogrel for preventing clot formation in patients with cardiovascular problems [1]), it also increases the risk of unexpected DDIs and adverse side effects [2]. Consequently, regulatory and monitoring institutions have developed various *in silico, in vitro, preclinical*, and *clinical* methods to identify potential DDIs and inform patients about their associated risks [3].

Monotherapy is often inadequate for effective treatment, as managing complex diseases typically requires drug combinations. Consequently, timely and costly clinical trials are necessary to ensure minimal side effects, a key focus of DDI research. Deep learning methods offer the potential to reduce these expenses through computational prediction of DDIs [4, 5].

Due to their pharmacological and pharmacokinetic properties, small-molecule drugs play a critical role in the treatment of diseases and the discovery of new drugs. Advances in deep learning methods have significantly accelerated the discovery of new small-molecule drugs, which now account for approximately 98% of all drugs [6]. As a result, DDI prediction for small-molecule drugs has also gained considerable attention.

A plethora of research studies have utilized computational methods for DDI prediction. HDN-DDI [7], SSF-DDI [8], DDI- Transform [9], BINDTI [10] and SumGNN [11] were GNN-based methods that used various graph neural network architectures to model molecular structures and interactions. MCNN- DDI [12], CNN-DDI [13] and CNN-Siam [14] utilized CNN-based architectures to extract features from drug sequences, proteins, or other drug-related data such as SMILES and targets.

Although most studies have focused on small-molecule drugs, it is essential not to overlook biotech drugs in health and medicine. Biotech drugs have proven highly effective in treating diseases with high morbidity and mortality, such as cancer [15]. For example, insulin information, a biotech drug, is vital for individuals with diabetes [16]. Therefore, it is crucial to emphasize that both small-molecule and biotech drugs warrant further investigation and exploration in drug discovery research. Small-molecule drugs have simple, well-defined chemical structures that can be represented using SMILES notation, making them easier to synthesize and modify. In contrast, biotech drugs are larger, protein-based molecules with complex structures that cannot be easily represented by SMILES. For example, Aspirin (Figure 1 in *Supplementary*) is a small-molecule drug with a well-defined SMILES representation, whereas Insulin Lispro (Figure 2 in *Supplementary*) is a biotech drug with a complex amino acid sequence that cannot be easily represented by SMILES [17]. These two types of drugs differ significantly in their physicochemical and pharmacokinetic properties. Additionally, they exhibit distinct behaviors in absorption, distribution, metabolism, and excretion within the body [18]. As a result, studying the interactions between these two types of drugs is of paramount importance in advancing drug discovery.

**Fig. 1.**
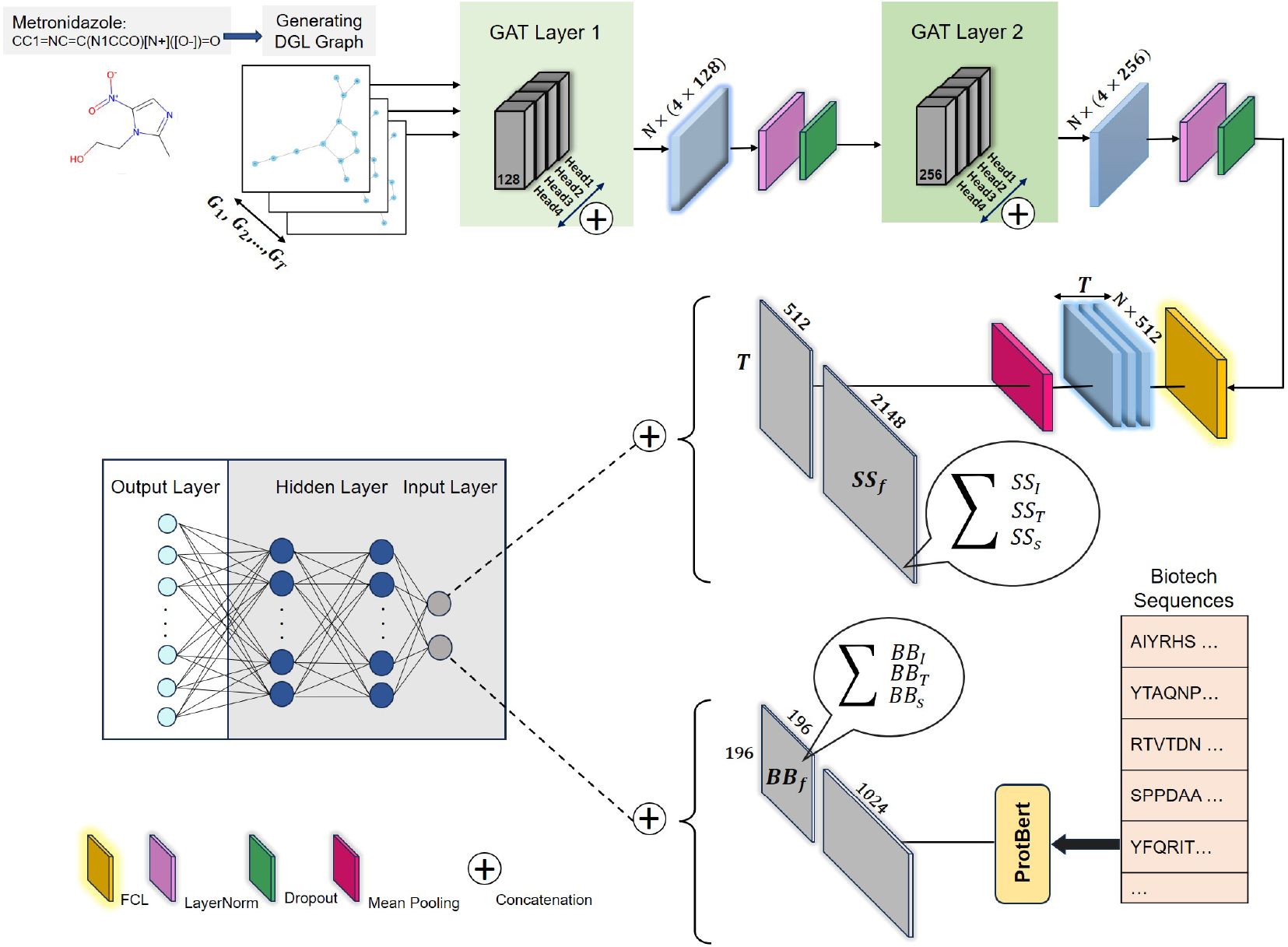
The BSI-Net framework predicts DDIs using small-molecule and biotech drug properties. For small-molecules, it converts SMILES strings to graphs using DGL, applies two GAT layers to generate representations, and concatenates these with similarity information (sum of *SS*_*I*_, *SS*_*T*_, *SS*_*S*_). For biotech drugs, it uses ProtBert to create sequence representations, which are then concatenated with similarity features. Finally, BSI-Net combines the small-molecule and biotech drug representations and inputs them to an MLP to predict interaction types.

**Fig. 2.**
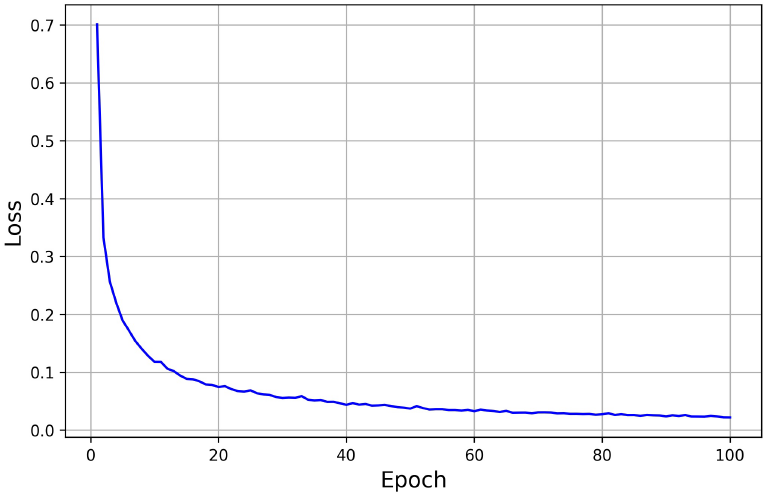
BSI-Net convergence.

This work proposes a graph-attention-based method with deep models that utilizes the similarity and sequence properties as well as the domain-specific language model methods of small-molecules and biotech drugs to predict their interactions. To do this, we generate and propose a new dataset that contains the last approved drugs from the DrugBank, with their related and significant relation with DDI prediction.

The structure of this paper is as follows. Section 2 introduces the related work briefly. Then, Section 3 introduces the generated data proposed in this work. Section 4 proposes BSI-Net method for biotech–small-molecule DDI prediction. Section 5 reports the results and finally, Section 6 concludes the paper.

## Related work

As mentioned earlier, there exists a plethora of research studies focused on small-molecule DDI prediction. Some of them has been briefly introduced in the *Supplementary* information. However, only a limited number of studies propose methods for biotech–small-molecule pair DDI prediction. We briefly describe these type of biotech-small molecules DDI predictions studies below.

Huang et al. [19] proposed a three-step method for predicting interactions between small-molecules and biotech drugs. In the first step, they utilized different feature representations of small-molecules and biotech drugs (e.g., structural and interactive properties) and applied operations such as Jaccard similarity computation and PCA to generate feature vectors. However, as our assessment and confirmation in [20] indicate, the authors did not compute one-hot encodings correctly and failed to perform the Jaccard computation accurately. They applied PCA directly on the incorrect one-hot encodings. Additionally, they employed a proposed type of negative sampling. The final step involved using a deep learning model to predict the types of interactions. However, they reported performance based on a dichotomization of negative and positive data and did not provide results for the correct prediction of interaction types.

Ru et al. [20] utilized a two-level graph representation: one level for capturing DDIs and DTIs and another for graph entities corresponding to each small-molecule drug. They applied a type of graph convolutional network to generate embeddings and subsequently used a multi-layer perceptron to predict interaction types between biotech–small-molecule pairs. However, they reported results primarily for small-molecule DDIs, which does not directly align with their main objective. Some biotech drugs have multiple sequences listed in DrugBank. Often, the first sequence is identical across several drugs, while variations occur in the subsequent sequences. For example, Insulin aspart, Insulin detemir, and Insulin glulisine all share the same first sequence, but differ in the second sequence. Based on the assessment of [19, 20], both studies consider only the first sequence for downstream tasks. In contrast, this work utilizes all sequences of a drug for downstream tasks.

In another study, [21], the authors explored machine and deep learning methods for predicting biotech–small-molecule DDIs. They applied machine learning to structural and interaction properties of the drugs to perform binary prediction. Their results indicated that CNN achieved the best performance for binary prediction. However, this work demonstrates that CNN is not the optimal choice for multiclass prediction. Furthermore, this work distinguishes itself in two key ways: (1) the introduction of a new dataset and (2) the proposal of a novel method.

## Dataset

To propose a new method for prediction of the interaction between biotech type drugs and small-molecule type drugs, we explore the literature. Huang et. al [19] proposed a dataset that contains information of those two types of drugs. However, The number of biotech was not enough for further analysis. Ru et. al [20] proposed two different datasets one for prediction bt-sm interactions and the other is just for small-molecule DDI predictions. The authors mentioned the problems of the former method and dataset. However, their dataset suffers from the same problem as the former. Their dataset has only the information of 55 biotech drugs. Although, this work has the aim of proposing a new learning model to improve the prediction performance, it introduces a new generated dataset of information of these two types of drugs. Thus, this section provides a description of the proposed dataset.

All the requirements for dataset generation was collected from DrugBank [17]. The dataset contains information of three identities, i.e., Biotech drugs (protein-based), small-molecule drugs (chemical drugs), proteins or targets that the drugs interacts with them. While the total number of reported biotech drugs in the DrugBank are 1620, but those drugs of this type which has been approved and their corresponding sequences are available is 196. Therefore, we collected the available info of this number of biotech drugs. Furthermore, the number of small-molecule drugs that are approved and their SMILES are available are equal to 2148. Thus, we collected their name and SMILES from the DrugBank. The total number of targets are 2186. But the unique number of total drugs (both types) that have an interaction with any protein is 1953. In other words, there exist drugs with no interaction with any target.

It is worth mentioning that there are two general types of interactions – drug-drug interactions (DDI) and drug-target interactions (DTI). DDIs itself are divided into three subtypes of biotech-biotech interactions (BBI), small-molecule - small-molecule interactions (SSI), biotech–small-molecule interactions (BSI). This work considers BSIs as the labels and aims to predict their values efficiently and effectively. There are 5034, 612947, 45205 number of BBIs, SSIs, and BSIs, respectively, therefore, totally 663186 interactions. the total number of DTIs are 7814. DTIs are divided into two types of biotech-target interactions (BTI) and small-molecule - target interactions (STI). There are 409 and 7405 of BTIs and STIs, respectively. The total number of unique targets that play a role in BTIs and STIs are 197 and 2056, respectively. Table 1 provides an overview of biotech and small-molecule drugs, including their targets and interactions.

**Table 1.**
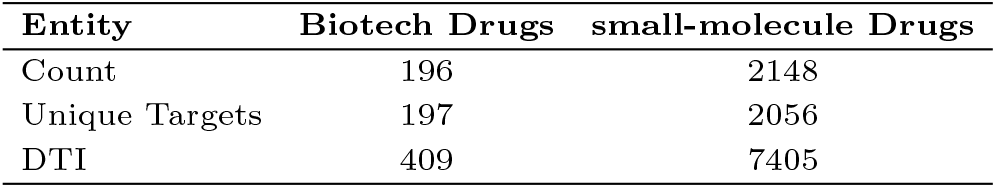
Statistics of drugs, targets, and interactions in the dataset.

| Entity | Biotech Drugs | small-molecule Drugs |
| --- | --- | --- |
| Count | 196 | 2148 |
| Unique Targets | 197 | 2056 |
| DTI | 409 | 7405 |

As mentioned above, the BSIs are considered as labels. We consider these labels as the positive labels which are equal to 45205. These positive labels are sentences that reports the interaction between each BSI pair. Figure 3 in *Supplementary* information represents some examples of such reports. We consider each of them as a separate label. There are 96 types of positive labels, with the frequencies of the labels ranging from 1 to 8809.

**Fig. 3.**
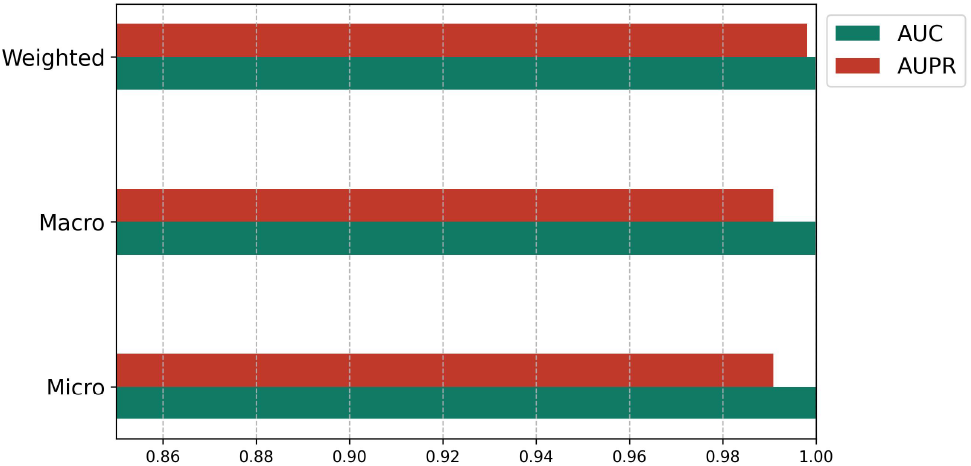
BSI-Net performance.

There are 31 BSI labels with frequencies above 100 (Table 1 in *Supplementary* information shows the statistics of the labels in descending order sorted). Therefore, we keep these as the number of positive types of labels, considering that the total number of existing BSIs is 43876. Having this primary raw information, we have prepared the dataset as follows.

- small-molecule drugs have SMILES and have interactions with targets. From these values, we computed four types of feature vectors for small-molecule drugs.
- We generate an interaction matrix for small-molecule drugs, resulting in a 2148 *×* 2148 matrix called “*SS*_*I*_ “.
- We used Tanimoto score [22] to compute the similarity among them. As mentioned earlier, there are 2148 drugs of type small-molecule in the dataset. Therefore, we use the similarity values to generate a feature vector for each drug with size 2148. We call this feature matrix “*SS*_*s*_”. Its dimension is 2148 *×* 2148.
- Then we generate the “*STI*” matrix (size 2148 × 2056), which represents the interactions between small-molecule drugs and targets. Then, we apply cosine similarity on the vectors within the “*STI*” matrix to calculate small-molecule similarities based on their interactions with the targets. The resulting matrix, called “*SS*_*T*_ “, has a size of 2148 *×* 2148.
- In addition to similarity checking, we generate bi- directed DGL graphs for each drug from its SMILES representation using DGL-LifeSci [23].
- The biotech drugs have sequences and known interactions with targets. From these, we derived multiple types of feature vectors.
- We generate an interaction matrix for biotech drugs, resulting in a 196 *×* 196 matrix called “*BB*_*I*_ “.
- We applied global sequence alignment [24] on the drugs’ sequences using a computational method in bioinformatics to calculate the similarity between the sequences of protein-based drugs. Global sequence alignment compares the entire length of two or more DNA, RNA, or protein sequences, aiming to identify structural and functional similarities between amino acids in the protein strands. It is important to note that some drugs have multiple sequences. For these drugs, in contrast with [19, 20],we concatenate the sequences and then apply the sequence alignment approach. Using the similarity values, we generate a feature vector for each drug, resulting in a similarity matrix with dimensions 196 *×* 196. We refer to this matrix as “*BB*_*s*_”.
- We compute a similarity matrix for biotech drugs based on their interactions with targets using cosine similarity. Notably, the set of 197 targets interacting with biotech drugs differs from those associated with small-molecules. Therefore, we construct the “*BTI*” matrix (size 196 *×* 197) to represent these interactions. Using cosine similarity, we then generate the biotech drug similarity matrix, called “*BB*_*T*_ “, with a size of 196 *×* 196.
- We utilized protein drug sequences and applied the ProtBert model to generate embedding vectors of size 1024 for these drugs. ProtBert is a pretrained language model based on the BERT architecture and Transformers, specifically designed for processing and analyzing protein sequences. By leveraging deep learning on a vast dataset of protein sequences, ProtBert can capture complex relationships between amino acids and extract biologically meaningful features from raw sequences [25]. For drugs with multiple sequence chains, we first computed separate embedding vectors for each chain and then averaged them to obtain a single embedding vector per drug. The resulting embeddings form a matrix called “*B*_*P rotB*_”, with dimensions 196 *×* 196.

Table 2 shows the summary of the generated features in the proposed dataset.

**Table 2.** Overview of Feature Representations in the Proposed Dataset.

| Drug Type | Feature Representation | Dimensions |
| --- | --- | --- |
| Small-molecule | Drug-drug interaction matrix ($SS_I$) | $2148 \times 2148$ |
| | Structural similarity matrix (Tanimoto, $SS_s$) | $2148 \times 2148$ |
| | Interaction-derived similarity matrix ($SS_T$) | $2148 \times 2148$ |
|  | Graph-based representation (DGL) | – |
| Biotech | Drug-drug interaction matrix ($BB_I$) | $196 \times 196$ |
| | Sequence similarity matrix (Global alignment, $BB_s$) | $196 \times 196$ |
| | Interaction-derived similarity matrix ($BB_T$) | $196 \times 196$ |
| | Protein sequence embeddings (ProtBert) | $196 \times 1024$ |

## Proposed Method

This section proposes BSI-Net, a new graph attention-based method for BSI predictions. Before delving into the proposed method, we mention the inputs and basis methods that we have examined in order to compare the BSI-Net performance.

From the BSI-data, we construct feature vectors for small-molecules and biotech drugs as follows. We use *SS*_*I*_, *SS*_*T*_, and *SS*_*s*_ for small-molecule feature vectors in such a way that the input is their summation *SS*_*f*_ = *SS*_*I*_ + *SS*_*T*_ + *SS*_*s*_, and for biotech drugs we use the summations of *BB*_*I*_, *BB*_*T*_, and *BB*_*s*_, i.e., *BB*_*f*_ = *BB*_*I*_ + *BB*_*T*_ + *BB*_*s*_ as their input vectors. To predict the interaction type between the *i*-th small-molecule and *j*-th biotech drug, we concatenate the *i*-th row of *SS*_*f*_ with the *j*-th row of *BB*_*f*_, forming the input vector for classification. We evaluate our approach using three machine learning models including Support Vector Machine (SVM), Random Forest (RF), and XGBoost. SVM is a powerful machine learning model useful in classification and regression that looks for a hyperplane with the maximum distance from the samples of the two categories [26]. The random forest is another high-performance machine learning model that uses a set of decision trees, each a subsample of features. Each tree decides whether the category is the input to the prediction models. The final choice for the random forest is the majority vote, which conceptually is based on the use of several weaker learning models to avoid overfitting in the final learning [27]. XGBoost is a fast and highly efficient machine learning algorithm that is based on gradient boosting, which in each iterate creates a new tree to reduce the construction errors of the previous tree. Furthermore, it assigns a larger weight to the samples that caused the error in the previous trees. This property leads to the XGBoost learning the model more efficiently. In the final step, all the trees are combined to produce a powerful learning model [28].

### Graph attention netwrok (GAT)

BSI-Net utilizes a graph attention network to have more significant embeddings. Thus, we describe the attention model before describing the proposed method. Given a graph *G* = (*V, E*), where *V* is the set of nodes and *E* is the set of edges, the graph attention network (GAT) updates the node embeddings using attention mechanisms [29]. The first step is feature transformation, in which, each node *i* has an input feature vector **h**_*i*_. The GAT first applies a linear transformation as follows.

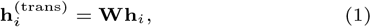

where **W** is a learnable weight matrix. Then, for each edge between nodes *i* and *j*, we calculate an attention score using the following equation.

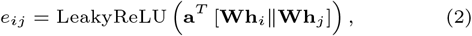

where **a** is a learnable attention vector, ∥ denotes concatenation, LeakyReLU is the activation function. Afterwards, the attention coefficients are computed. In other words, attention scores are normalized using the softmax function as follows.

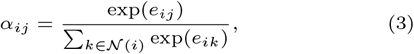

where 𝒩 (*i*) is the set of neighbors of node *i*. The new representation of node *i* is calculated by aggregation of the feature vectors of the its neighbors.

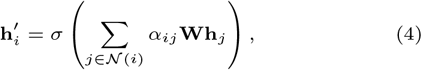

where *σ* is a non-linear activation function, e.g., ReLU. For stability and efficient learning, multiple attention heads are used, as follows.

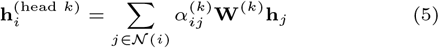

The final node representation is computed by either averaging,

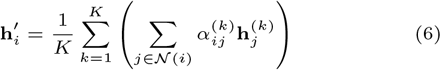

or concatenating the outputs of multiple heads.

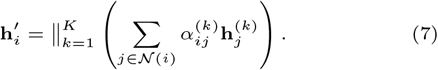

BSI-Net concatenates the outputs of multiple attention heads to form the final node representation.

### BSI-Net

We have introduced the main components of our proposal, BSI-Net. In this section, we will describe the details of the framework. Figure 1 illustrates the general structure of BSI-Net. The inputs are categorized into two types: small-molecule drug properties and biotech drug properties.

For small-molecule properties, we categorize and process them in two ways. The first type of property is derived from the SMILES representation of the drugs. BSI-Net converts these SMILES into graphs using DGL (Deep Graph Library). It then applies a multi-head graph attention layer with a size of 128 to each graph, utilizing four multi-head attention mechanisms in this layer. The output for each node in the graph (N nodes in total) results in a matrix of size 4*×*128. Following this, BSI-Net applies normalization and a dropout layer with a drop rate of 0.2 to the node features. A second multi-head graph attention layer, this time with a size of 256, is then applied, followed by another normalization layer and dropout layer. The processed data is then fed into a fully connected layer. Finally, a mean pooling layer is applied, resulting in feature vectors of size 512 for the small-molecule drugs.

The additional properties of the small-molecules are derived from similarity matrices. The summed results of *SS*_*I*_, *SS*_*T*_, and *SS*_*S*_ yield *SS*_*f*_ = *SS*_*I*_ + *SS*_*T*_ + *SS*_*S*_, which constitutes the next component of the small-molecule feature vectors, with a feature size of 2148. The output from the graph attention layer is concatenated with *SS*_*f*_, giving a final feature vector size of 2660 for small-molecule drugs.

The biotech properties are further divided into two subsets. One subset is derived from the summation of the similarity matrices: *BB*_*f*_ = *BB*_*I*_ + *BB*_*T*_ + *BB*_*S*_ with a feature size of 196. Additionally, the biotech sequences are processed through the pre-trained ProtBERT, resulting in feature vectors of size 1024. The final feature vectors for biotech drugs are produced by concatenating the results from ProtBERT and *BB*_*f*_, leading to feature vectors with a size of 1220.

The previous steps lead to generating embeddings for small-molecules and biotech drugs. The final feature vector for DDI prediction is the concatenation of each small-molecule drug feature vector with the biotech feature vector. These final vectors are the inputs for the prediction part. BSI-Net uses an MLP for multiclass prediction. The network contains two hidden layers. The input layer has a size of 3880, which consists of concatenated embeddings from the small-molecule and biotech drug feature vectors. The output layer contains 32 neurons, corresponding to the number of interaction classes. The activation function in the hidden layers is ReLU, while the output layer uses the softmax function to predict the interaction class probabilities. BSI-Net employs the cross-entropy loss function to compute the prediction error, as defined in Equation (8). The model is optimized using the Adam optimizer, and hyperparameter tuning is performed via grid search with various configurations of hidden layer sizes, dropout rates, and learning rates.

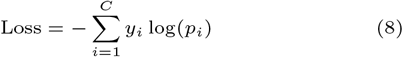

The steps for performing BSI-Net are outlined in Algorithm 1. The algorithm requires several inputs: SMILES, the feature set of small-molecules (*SS*_*f*_), sequences of biotech drugs (*BB*_*f*_), labels (Y), and an error threshold (*ϵ*). The input data is divided into multiple folds. In each fold, the training phase continues as long as the training error exceeds the predefined threshold, as detailed in lines 4 to 14.

During the training phase, the SMILES data from the training subset (*SMILES*^(*\k*)^) are processed using DGL to create small-molecule graphs, as shown in line 5. Line 6 applies the proposed Graph Attention Network (GAT) model to the output from DGL. The algorithm then concatenates the feature set of the training subset, 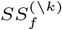, resulting in a feature vector for the small-molecules, denoted as *S*_*T*_.

In lines 8 and 9, the feature vectors for biotech drugs are prepared. Line 10 concatenates the feature vectors of small-molecules and biotech drugs. Line 11 applies a Multi-Layer Perceptron (MLP) model to the combined feature vector to evaluate the interactions among drugs during the training phase. Line 12 calculates the loss for the predicted training values, following the loss function defined in Eq. (8). Based on this calculated error, the network weights are updated through backpropagation in line 13.

Once the training subset converges, the labels for the test subset are computed. This process is repeated for all folds. Finally, the evaluation results are compiled in line 19.

**Algorithm 1.**
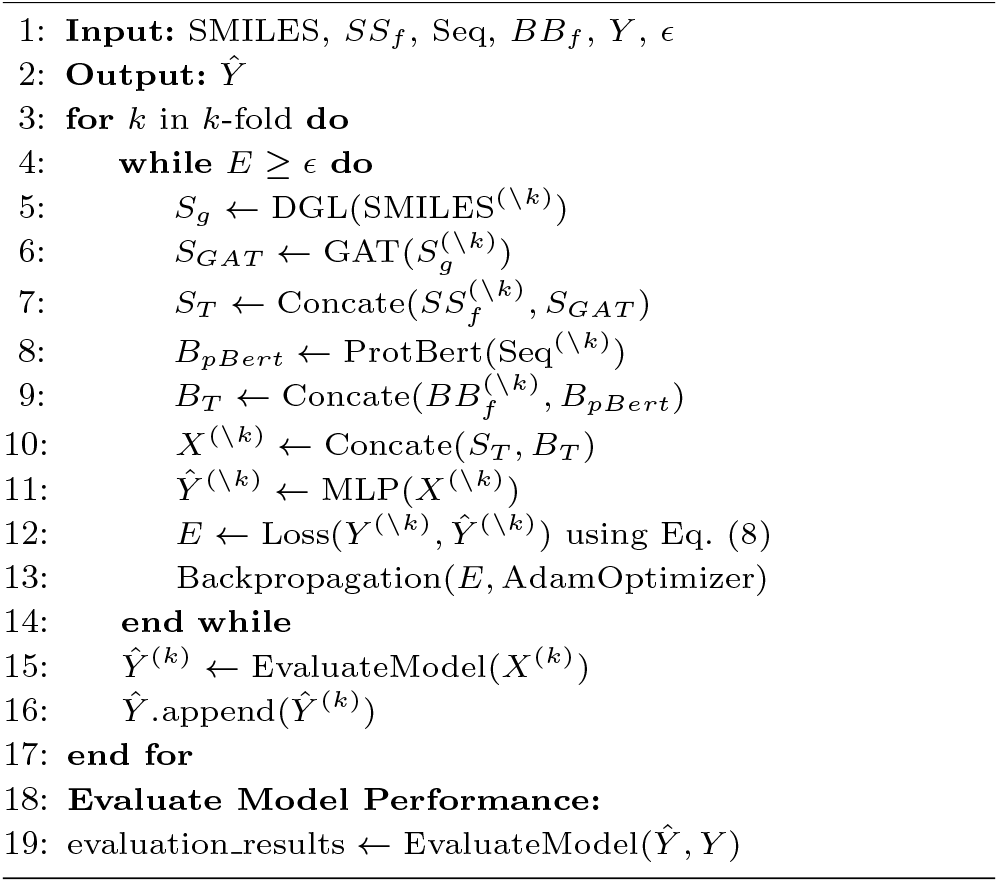
BSI-Net

## Results

We employ a k-fold stratified cross-validation approach to evaluate the performance of BSI-Net. The stratified division ensures that each fold maintains the same label distribution as the overall dataset, which is particularly important for handling imbalanced labels. Consequently, both the training and test sets contain a proportional representation of each label [30].

As mentioned earlier, our dataset consists of 31 positive labels and one negative label, totaling 32 labels. To assess model performance comprehensively, we report results based on three categories of evaluation metrics: micro, macro, and weighted. These metrics are particularly useful when dealing with multi-class classification tasks [31].

We report the evaluation metrics for method comparison in three different regime of micro, macro, and weighted. Their full description is in Section 5 of the *Supplementary* information.

To ensure a comprehensive performance evaluation, we compare the results of the mentioned random forest, SVM, and XGBoost models. Additionally, we implemented a three-layer multilayer perceptron (MLP) for comparison. Furthermore, we employ the CNN architecture introduced in [32, 33], which consists of three layers. Each layer includes a convolutional layer, batch normalization, and a dropout mechanism. Following these layers, the architecture incorporates a dense layer and concludes with an activation layer. Finally, we compare the performance of BSI-Net with the results reported in [19, 20]. Figure 4 in *Supplementary* information shows the confusion matrix of the predicted labels against the true labels of 32 labels (31 positive labels plus the negative label).

### Micro

We have introduced micro results to evaluate the performance of the proposed method. Tables 3 and 4 report the results based on micro performance. The former provides accuracy, precision, recall, and F1-score; the latter presents MCC, AUROC, and AUPR. Regarding micro accuracy, BSI-Net achieves the highest performance, followed by the machine learning approaches, particularly random forest, which ranks next; the MLP method follows, while the CNN model ranks last with the lowest performance. Notably, machine learning approaches demonstrate high performance with lower complexity compared to the MLP and CNN models. The same pattern is observed for precision, recall, and F1-score in Table 3. Table 4 shows a similar pattern to the previous table. BSI-Net exhibits the best performance except in AUPR, where it ranks second, and random forest takes the top position. Again, this underscores the necessity for researchers to consider the strength of machine learning methods when proposing techniques. It is worth mentioning that the values in each row of Table 3 are equal. Therefore, the authors verified that all results are correct, and this observation is typical in micro value computations, making it less favorable for evaluating performance.

**Table 3.** Evaluation of Methods in Micro Regime(first part)

| Methods | Accuracy(std) | Precision(std) | Recall(std) | F1-Score(std) |
| --- | --- | --- | --- | --- |
| **SVM** | 0.9764(0.001) | 0.9764(0.001) | 0.9764(0.001) | 0.9764(0.001) |
| **RF** | 0.9827(0.001) | 0.9827(0.001) | 0.9827(0.001) | 0.9827(0.001) |
| **XGBoost** | 0.9730(0.002) | 0.9730(0.002) | 0.9730(0.002) | 0.9730(0.002) |
| **MLP** | 0.7381(0.014) | 0.7381(0.014) | 0.7381(0.014) | 0.7381(0.014) |
| **CNN** | 0.6048(0.048) | 0.6048(0.048) | 0.6048(0.048) | 0.6048(0.048) |
| **BSI-Net** | **0.9861(0.009)** | **0.9861(0.009)** | **0.9861(0.009)** | **0.9861(0.009)** |

**Table 4.** Evaluation of Methods in Micro regime (second part).

| Methods | MCC(std) | AUROC(std) | AUPR(std) |
| --- | --- | --- | --- |
| **SVM** | 0.9675(0.002)) | 0.9997(0.000) | 0.9946(0.000) |
| **RF** | 0.9761(0.002) | 0.9997(0.000) | **0.9974(0.000)** |
| **XGBoost** | 0.9625(0.002) | 0.9998(0.000) | 0.9953(0.001) |
| **MLP** | 0.6941(0.013) | 0.9877(0.001) | 0.7980(0.014) |
| **CNN** | 0.5629(0.040) | 0.9781(0.004) | 0.6621(0.053) |
| **BSI-Net** | **0.9809(0.012)** | **1.0000(0.000)** | 0.9908(0.009) |

### Weighted

In addition to micro and macro averages, weighted evaluation metrics were computed (Table 5). BSI-Net performed best, followed by random forest, while CNN performed worst. Considering all three evaluation modes (micro, macro, and weighted), BSI-Net achieved the highest performance, followed by XGBoost and random forest. MLP and CNN exhibited the lowest performance, with MLP outperforming CNN. Furthermore, the results for the macro regime has been reported in *Supplementary* information.

**Table 5.** Evaluation of Methods in Weighted Regime.

| Methods | Precision (std) | Recall (std) | F1-Score (std) | AUROC (std) | AUPR (std) |
| --- | --- | --- | --- | --- | --- |
| **SVM** | 0.9770(0.001) | 0.9764(0.001) | 0.9765(0.001) | 0.9986(0.000) | 0.9906(0.001) |
| **RF** | 0.9826(0.001) | 0.9827(0.001) | 0.9824(0.001) | 0.9995(0.000) | 0.9961(0.000) |
| **XGBoost** | 0.9729(0.002) | 0.9730(0.002) | 0.9724(0.002) | 0.9991(0.000) | 0.9935(0.001) |
| **MLP** | 0.8592(0.007) | 0.7381(0.014) | 0.7655(0.014) | 0.9788(0.002) | 0.9294(0.006) |
| **CNN** | 0.7981(0.013) | 0.6048(0.048) | 0.6367(0.047) | 0.9473(0.009) | 0.8540(0.014) |
| **BSI-Net** | **0.9866(0.022)** | **0.9861(0.025)** | **0.9861(0.021)** | **0.9999(0.000)** | **0.9981(0.007)** |

Table 6 compares BSI-Net’s performance against state-of-the-art methods (Multi-SBI [19] and CB-TIP [20]), but a direct comparison is limited due to varying numbers and types of drugs used, detailed in the table. BSI-Net and CB-TIP [20] results are weighted, while Multi-SBI [19] reports binary (positive/negative) classification results instead of multiclass metrics. Furthermore, CB-TIP uses more small-molecules than BSI-Net, which exclusively considers approved drugs from the latest DrugBank list, unlike CB-TIP.

**Table 6.** BSI-Net outperforms state-of-the-art methods (Multi-SBI [19] and CB-TIP [20]) in weighted biotech–small-molecule DDI prediction. ^*†*^: Multi-SBI’s multiclass results were reported as a binary dichotomization, as they did not report a weighted version.

| Method | Biotech | Small-molecules | F1-score(std) | AUROC(std) | AUPRC(std) |
| --- | --- | --- | --- | --- | --- |
| Multi-SBI<sup>†</sup> | 148 | 1941 | 0.8673 | 0.9997 | 0.9892 |
| CB-TIP | 55 | 3119 | 0.9780 ± 0.08 | 0.9666 ± 0.05 | 0.9420 ± 0.10 |
| BSI-Net | 196 | 2148 | **0.9861 ± 0.021** | **0.9999 ± 0.000** | **0.9981 ± 0.007** |

Figure 2 shows BSI-Net convergence in 100 epochs. Figure 3 presents AUROC and AUPR bar charts for BSI-Net in three modes.

Table 5 in *Supplementary* information presents a subset of biotech–small-molecule drug pairs along with their corresponding current interaction labels and the predicted labels generated by BSI-Net.

## Conclusion

This work presents a novel DDI prediction method for biotech–small-molecule drug pairs, leveraging graph attention networks, domain-specific language model representations, and MLP analysis of drug properties. We introduce a new dataset and demonstrate the superior performance of our approach. Furthermore, we highlight the relevance of comparing our method with lighter machine learning models, which can offer sufficient performance for drug interaction prediction in drug discovery. Our method currently relies on similarity vectors of the drugs, and future research will explore using direct data representations for potentially improved results. Improved sampling enhances performance. CNNs, effective for binary prediction, struggle with multiclass problems. Exploring anomaly detection as an alternative to negative sampling is crucial for better interaction prediction. Table 2 and Table 3 in *Supplementary* information show best tuned hyperparameters of the implemented methods.

## Data Availability

The datasets and code generated in this study are available in the GitHub repository: https://github.com/BioinformaticsIASBS/BSI-Net.

Key Points

- Proposing a method that predicts small molecule- biotech drug pairs by integrating structural, sequential, and similarity features with graph attention and deep learning.
- Creating a dataset of approved biotech and small- molecule drugs from DrugBank, including available sequences or SMILES, to ensure all drugs are included for accurate BSI prediction.
- BSI-Net achieves superior performance compared to state-of-the-art methods across all evaluation metrics.

## Acknowledgments

The authors thank Javad Asghari for dataset generation and Masih Hajsaeedi for sequence alignment.

## Author contributions statement

M.H. and F.N. conceptualized the study. M.H. supervised and managed the project, leading the writing of the draft and revisions. F.N. contributed to method development, data curation, validation, visualization, and manuscript refinement.

