## Supplemental material for "A Graph-Attention-Based Deep Learning Network for Predicting Biotech–Small-Molecule Drug Interactions"

### 1 Small Molecule vs. Biotech

Figure 1 and Figure 2 demonstrate the structure of Aspirin as a Small molecule and Insulin Lispro as a Biotech drug [1].

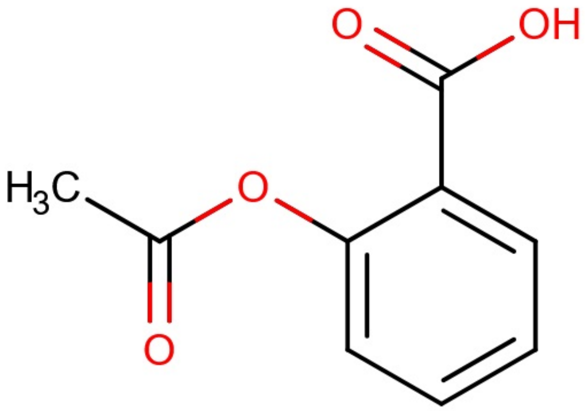


Figure 1: Structure of Aspirin, a small-molecule drug with its atomic arrangement and bonds.

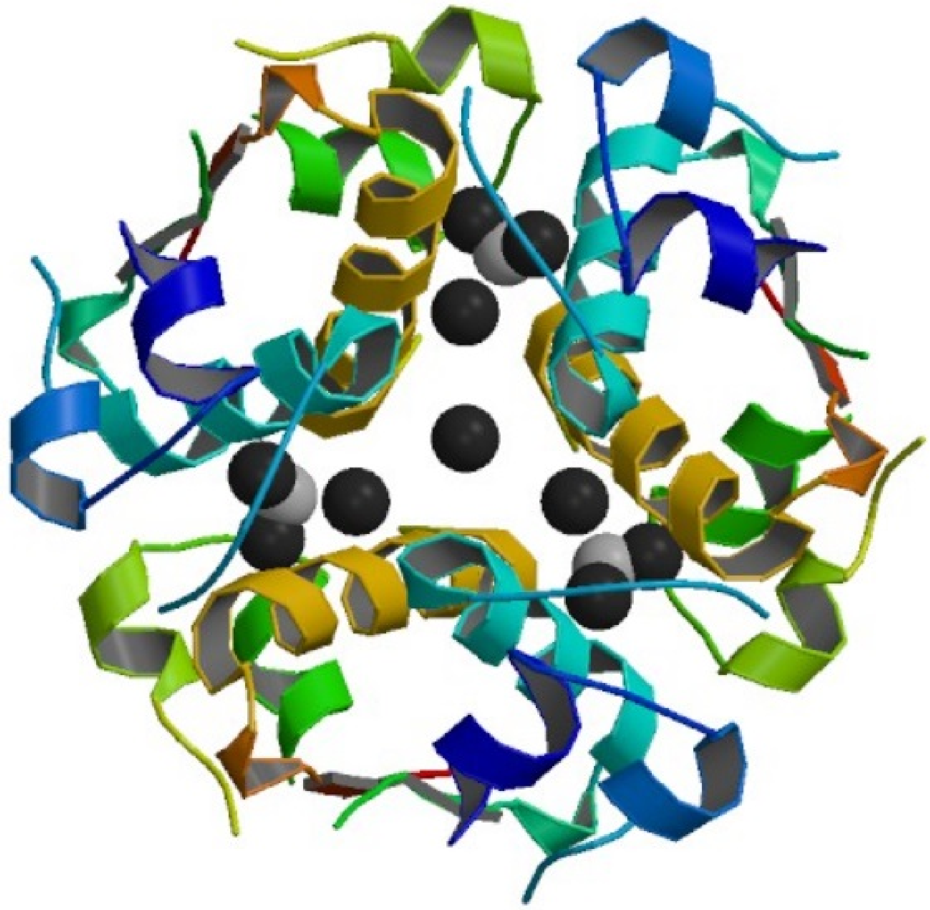

Figure 2: Structure of Insulin Lispro, a biotech drug with its modified protein sequence.

#### 2 Related work

In the field of drug-drug interaction prediction, machine learning methods can be divided into six categories: Similarity-based approaches, Traditional classification, Network diffusion, Matrix factorization, Ensemble-based approaches, and literature-based [2]. To predict conventional and synthetic DDIs, Yu et al. [3] developed a DDINMF approach based on semi-non-negative matrix factorization. Using drug features and known DDI information, Zhang et al. [4] proposed a prediction model combining neighbor-recommendation, random walk, and matrix perturbation strategies in a versatile ensemble framework.

Ryu et al. [5] introduced DeepDDI, a deep learning framework that predicts drug-drug and drug-food interactions using compound names and SMILES-based structural similarity profiles. The framework converts molecular structures into feature vectors and processes them through a multi-label neural network to identify possible interaction types. Yan et al. [6] introduced DDI-IS-SL, a method for predicting drug-drug interactions by combining different types of drug similarity. This method calculates similarities using chemical, biological, and phenotypic data, incorporates additional similarity measures using a Gaussian kernel, and then applies a machine learning model to predict interaction scores. Zhu et al. [7] introduced DGDFS, a feature selection method for predicting adverse DDIs. This method uses dependency-guided learning to select features based on their interdependencies, focusing on feature relationships within large datasets. Chen et al. [8] proposed MUFFIN, a multi-scale feature fusion framework for DDI prediction. The framework integrates features from different scales to capture complex relationships between drugs, utilizing both molecular and network-level information.

Lin et al. [9] developed MDF-SA-DDI, a deep learning model that predicts drug-drug interactions using multiple types of drug information based on similarity. The model employs neural networks to extract features from the data and utilizes a transformer architecture to integrate them for improved predictions. Yu et al. [10] proposed RANEDDI, a model for predicting DDIs using relation-aware network embedding. This approach captures both the structural features of drug networks and the specific types of relationships between drugs to generate more accurate interaction predictions. Al-Rabeah and Lakizadeh [11] proposed a graph neural network approach for predicting DDIs and their associated effects. Their model constructs a network representation of drugs from various data sources, generates drug embeddings, and employs a deep learning architecture to predict both the occurrence and type of DDI events. Zhong et al. [12] proposed DDI-GCN, a framework that uses graph convolutional networks (GCNs) to predict DDIs by analyzing drug chemical structures.

Zhu et al. used CNN and attention-based for sequence features and neural network, GAT, and SAGPooling for substructure features, combining them for

DDI prediction [13]. Asfand et al. employed a CNN model to predict DDIs. It integrated multiple drug features, including SMILES, enzymes, pathways, and targets, using four CNN sub-models, which are later concatenated for final prediction [14]. Su et al. used a neural network that predicts DDIs by combining drug structure and protein-binding features. It learns important patterns using graph-based learning [15]. Peng and colleagues proposed a bi-directional network that integrated multi-head attention and intention mechanisms to fuse drug and protein features. It first encodes drugs using GCN and proteins using a combination of CNN and self-attention [16]. Sun and colleagues utilized a GAT-based hierarchical dual-view representation learning network. Their network enables DDI prediction by capturing both local substructure interactions and global molecular information [17].

### 3 Biotech-small molecule DDI Labels

Figure 3 represents four samples of positive labels in our dataset.

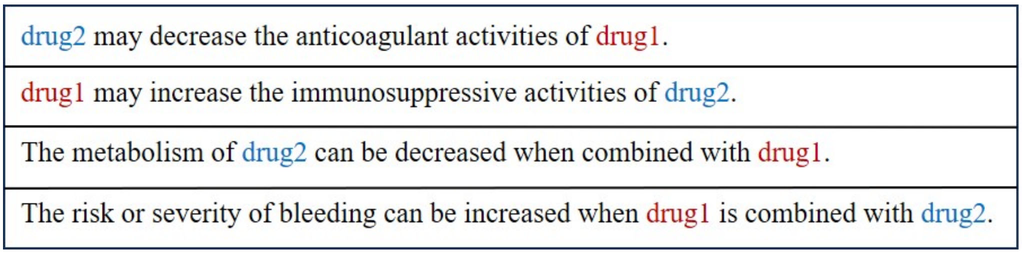


Figure 3: Four examples of different positive labels.

Table 1 demonstrates that there are 31 BSI labels with frequencies above 100. Therefore, we keep these as the number of positive types of labels, considering that the total number of existing BSIs is 43876.

Table 1: Distribution of the 96 positive labels used in the dataset, categorized by frequency range.

| **Count of labels** | **Frequency Range** |
|---|---|
| 3 | $> 4000$ |
| 7 | $> 2000$ |
| 10 | $> 1000$ |
| 15 | $> 500$ |
| 31 | $> 100$ |
| 41 | $> 50$ |

### 4 Parameter Tuning and Enhancements

To ensure robust and accurate classification performance, all models were individually fine-tuned through extensive hyperparameter exploration. For deep learning approaches—BSI-Net (based on a graph attention network), CNN, and multi-layer perceptron (MLP)—a range of configurations were tested, including variations in learning rate, dropout rate, activation functions, and optimization algorithms. The optimal settings were selected based on validation performance, with all three models achieving their best results using the Adam optimizer and ReLU activation function but differing in dropout rates and architectural structure. These final hyperparameter settings for the deep learning models are summarized in Table 2.

Table 2: Best hyperparameter settings for deep learning models.

| **BSI-Net** | **CNN** | **MLP** |
|---|---|---|
| LR: 0.001<br>Optimizer: Adam<br>Activation: ReLU<br>Dropout: 0.2<br>Output: Softmax | LR: 0.001<br>Optimizer: Adam<br>Activation: ReLU<br>Dropout: 0.5<br>Output: Softmax | LR: 0.001<br>Optimizer: Adam<br>Activation: ReLU<br>Dropout: 0.3<br>Output: Softmax |

Similarly, traditional machine learning models were tuned using grid search across relevant hyperparameters. The support vector machine (SVM) achieved optimal performance with a linear kernel and a regularization parameter of $C = 10$. The best configuration for the random forest (RF) classifier included 10 estimators, the `log_loss` splitting criterion, and the `log2` method for feature selection. XGBoost was fine-tuned by varying the number of estimators, tree depth, and sampling ratios, resulting in the optimal combination of 100 estimators, a maximum depth of 3, a learning rate of 0.1, and both subsample and colsample_bytree set to 0.5. These optimized hyperparameters for the classical models are detailed in Table 3, and were used consistently throughout the evaluation process.

Table 3: Best hyperparameter settings for traditional machine learning models.

| **SVM** | **RF** | **XGBoost** |
|---|---|---|
| Kernel: Linear<br>C: 10 | n_estimators: 10<br>Criterion: log_loss<br>Max Features: log2 | n_estimators: 100<br>Max Depth: 3<br>LR: 0.1<br>Subsample: 0.5<br>Colsample_bytree: 0.5 |

### 5 Metrics Analysis

The micro set of evaluation metrics treats all classes equally and focuses on the overall performance of the model. Furthermore, its preference is for the larger classes. The micro metrics, compares all true predictions (TP and TN) and all false predictions (FP and FN)of each class together. Therefore, it does not show the information at the class level. In other words, it does not care about how the model performs in individual classes. While Reporting micro metrics may show a high performance of the general model, it may have a very low performance for the classes with low frequency [18]. There are several evaluation metrics in this category and we choose some of them that are suitable for this work, i.e., accuracy, precision, recall, F1-Score, BACC, MCC, AUC and AUPR.

The macro set of evaluation metrics treats all classes equally, regardless of their size. Unlike micro-averaged metrics, macro metrics compute performance for each class separately based on its own samples and then average the results across all classes. Essentially, macro-evaluation means that the model's overall performance is the average of its performance on each class with any distribution. More precisely, the importance of each class does not depend on its size. For example, in medicine and disease diagnosis and prognosis, are considered just as important as common diseases. Furthermore, assigning equal weights to all classes ensures that the evaluation metrics are not dominated by larger classes [19]. For this study, we select several macro-based evaluation metrics, including precision, recall, F1 score, AUC, and AUPR.

The weighted category lies between micro and macro evaluation. While it evaluates each class separately, like macro-averaging, it does not treat all classes equally. Instead, it assigns a weight to each class based on its size, meaning larger classes have a greater influence on the final score. More precisely, the overall evaluation is a weighted sum of the metrics computed for each class [19]. For this study, we select several suitable weighted evaluation metrics, including precision, recall, F1 score, AUC, and AUPR.

### 6 Extra results

#### 6.1 Confusion Matrix

Figure 4 shows the confusion matrix of the predicted labels against the true labels of 32 labels ( 31 positive labels plus the negative label). As the figure shows, most of the labels are predicted correctly. This confusion matrix is the basis for the following results.

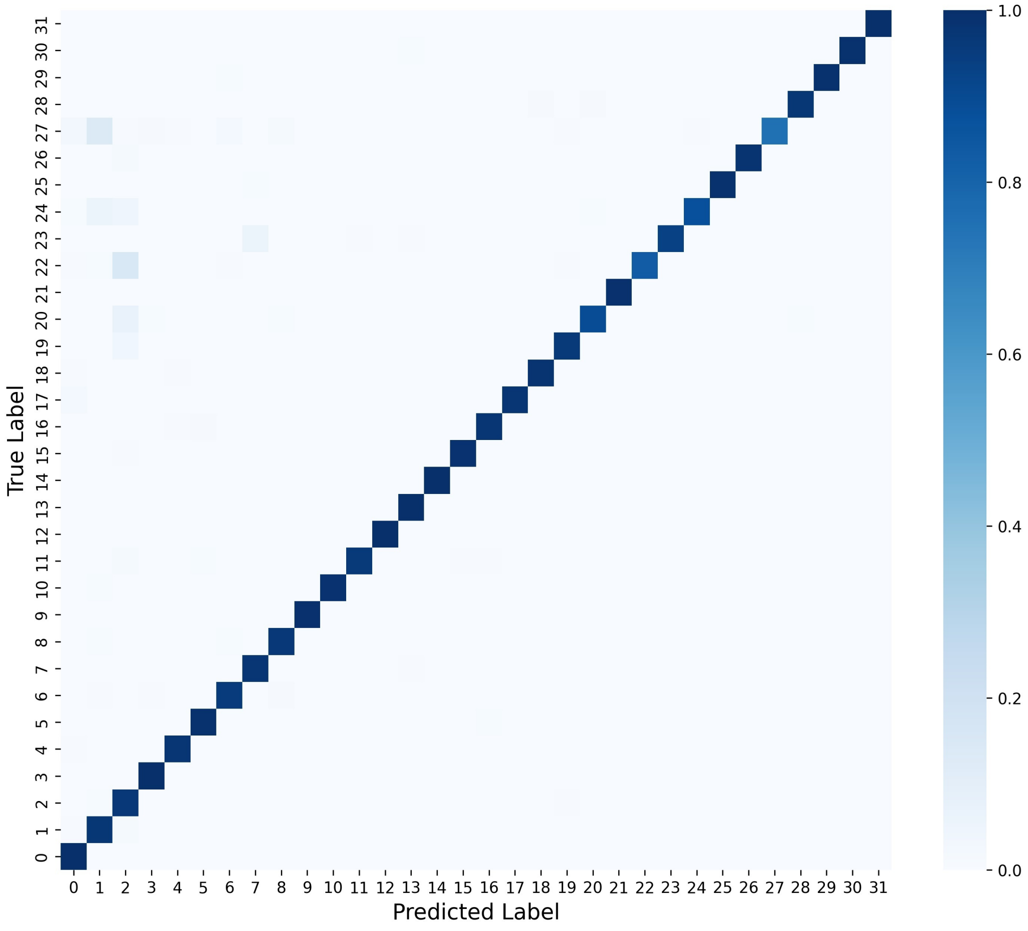


Figure 4: Confusion matrix for biotech–small-molecule DDI prediction.

#### 6.2 Macro

We discussed the issues with reporting the evaluation metrics in the micro regime. Now, we will report the results in the macro regime. Table 4 presents the evaluation metrics in the macro regime. BSI-Net shows the best performance across all metrics, while XGBoost takes the next rank. CNN has the lowest performance; in fact, all models except CNN achieve AUPR values above $0.9$.

Table 4: Evaluation of Methods in Macro Regime

| **Methods** | **Precision(std)** | **Recall(std)** | **F1-Score(std)** | **AUROC(std)** | **AUPR(std)** |
|---|---|---|---|---|---|
| **SVM** | 0.9649(0.002) | 0.9562(0.006) | 0.9598(0.003) | 0.9989(0.000) | 0.9783(0.004) |
| **RF** | 0.9775(0.004) | 0.9510(0.006) | 0.9614(0.006) | 0.9986(0.001) | 0.9831(0.005) |
| **XGBoost** | 0.9721(0.004) | 0.9410(0.003) | 0.9532(0.002) | 0.9990(0.000) | 0.9837(0.002) |
| **MLP** | 0.5632(0.020) | 0.9228(0.004) | 0.6551(0.018) | 0.9933(0.001) | 0.9040(0.005) |
| **CNN** | 0.4677(0.026) | 0.8619(0.010) | 0.5476(0.029) | 0.9855(0.002) | 0.8007(0.014) |
| **BSI-Net** | **0.9705(0.053)** | **0.9609(0.073)** | **0.9627(0.058)** | **0.9999(0.000)** | **0.9908(0.026)** |

##### 6.3 Comparison

Figure 5 compares BSI-Net's AUROC and AUPR in weighted mode with state-of-the-art methods, though this comparison is not entirely fair due to dataset differences. Multi-SBI [20] only reported binary results, but BSI-Net outperforms all compared methods, with random forest performing second best.

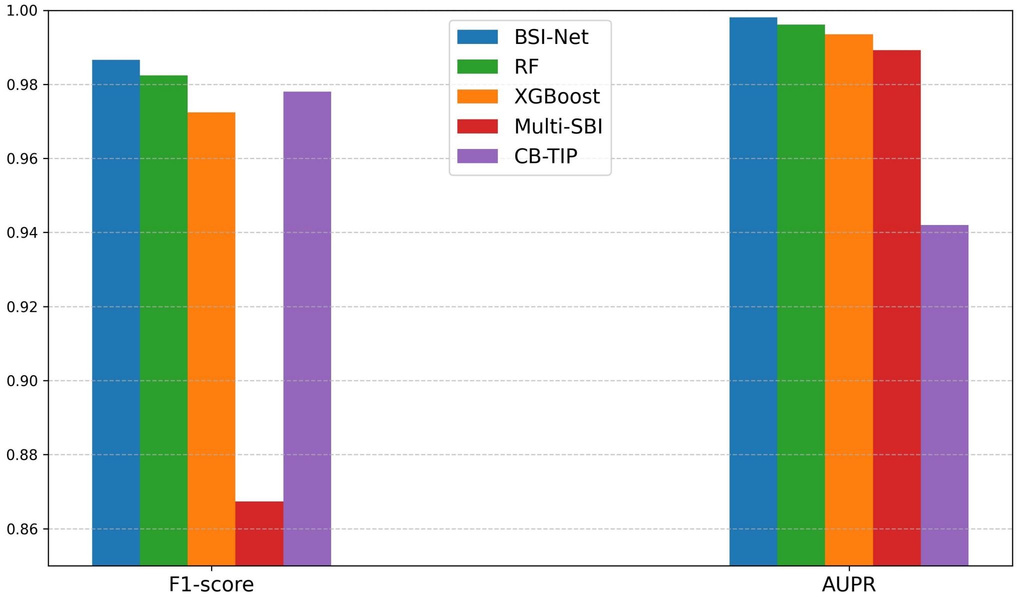


Figure 5: Comparison of BSI-Net with other methods based on F1-score and AUPR.

### 7 Predicted Labels

Table 5 presents a subset of biotech–small-molecule drug pairs along with their corresponding current interaction labels and the predicted labels generated by BSI-Net.

Table 5: Biotech–Small-molecule Drug Pairs with True and Predicted Labels

| **Biotech Drug** | **Small-molecule Drug** | **True Label** | **Predicted Label** |
|---|---|---|---|
| Insulin_human | Praziquantel | 0 | 1 |
| Darbepoetin_alfa | Cilazapril | 0 | 27 |
| Erythropoietin | Finafloxacin | 0 | 17 |
| Darbepoetin_alfa | Perindopril | 0 | 27 |
| Insulin_human | Trofinetide | 0 | 5 |
| Insulin_human | Diclofenac | 0 | 1 |
| Darbepoetin_alfa | Enalaprilat | 0 | 27 |
| Insulin_human | Pirfenidone | 0 | 1 |
| Darbepoetin_alfa | Trandolapril | 0 | 27 |
| Anakinra | Fosnetupitant | 0 | 1 |
| Insulin_human | Primaquine | 0 | 1 |
| Darbepoetin_alfa | Quinapril | 0 | 27 |
| Anakinra | Tivozanib | 0 | 1 |
| Darbepoetin_alfa | Cycloguanil | 0 | 17 |
| Darbepoetin_alfa | Moexipril | 0 | 27 |
| Erythropoietin | Cinnamaldehyde | 0 | 17 |
| Darbepoetin_alfa | Chlortetracycline | 0 | 17 |
| Darbepoetin_alfa | Ramipril | 0 | 27 |
| Insulin_human | Leflunomide | 0 | 1 |
| Darbepoetin_alfa | Spirapril | 0 | 27 |
| Sargramostim | Topotecan | 0 | 24 |
| Anakinra | Gepirone | 0 | 1 |
| Darbepoetin_alfa | Tafenoquine | 0 | 17 |
| Insulin_human | Binimetinib | 0 | 1 |
| Insulin_human | Aliskiren | 0 | 5 |
| Anakinra | Bezafibrate | 0 | 1 |
| Darbepoetin_alfa | Fosinopril | 0 | 27 |
| Insulin_human | Levobupivacaine | 0 | 1 |
| Darbepoetin_alfa | Artemotil | 0 | 17 |
| Insulin_human | Clopidogrel | 0 | 1 |
| Darbepoetin_alfa | Azithromycin | 0 | 17 |
| Urokinase | Nitroglycerin | 0 | 4 |
| Anistreplase | Nitroglycerin | 0 | 4 |
| Insulin_human | Zileuton | 0 | 1 |
| Erythropoietin | Chlortetracycline | 0 | 17 |
| Interferon_alfa-n1 | Telbivudine | 0 | 2 |
| Darbepoetin_alfa | Rescinnamine | 0 | 27 |
| Darbepoetin_alfa | Lefamulin | 0 | 17 |
| Insulin_human | Dacarbazine | 0 | 1 |
| Interferon_alfa-n1 | Dosulepin | 0 | 8 |
| Anakinra | Prednisolone_phosphate | 0 | 1 |
| Etanercept | Levomilnacipran | 0 | 1 |
| Anakinra | Lumacaftor | 0 | 1 |
| Erythropoietin | Chloramphenicol | 0 | 17 |

[16] Lihong Peng, Xin Liu, Long Yang, Longlong Liu, Zongzheng Bai, Min Chen, Xu Lu, and Libo Nie. Bindti: a bi-directional intention network for drug-target interaction identification based on attention mechanisms. *IEEE Journal of Biomedical and Health Informatics*, 2024.

[17] Jinchen Sun and Haoran Zheng. Hdn-ddi: a novel framework for predicting drug-drug interactions using hierarchical molecular graphs and enhanced dual-view representation learning. *BMC bioinformatics*, 26(1):28, 2025.

[18] Marina Sokolova and Guy Lapalme. A systematic analysis of performance measures for classification tasks. *Information processing & management*, 45(4):427–437, 2009.

[19] Maria Cristina Hinojosa Lee, Johan Braet, and Johan Springael. Performance metrics for multilabel emotion classification: comparing micro, macro, and weighted f1-scores. *Applied Sciences*, 14(21):9863, 2024.

[20] Dingkai Huang, Hongjian He, Jiaming Ouyang, Chang Zhao, Xin Dong, and Jiang Xie. Small molecule drug and biotech drug interaction prediction based on multi-modal representation learning. *BMC bioinformatics*, 23(1):561, 2022.
